# Functional analysis of the archaea, bacteria, and viruses from a halite endolithic microbial community

**DOI:** 10.1101/029934

**Authors:** Alexander Crits-Christoph, Diego R. Gelsinger, Bing Ma, Jacek Wierzchos, Jacques Ravel, Alfonso Davila, M. Cristina Casero, Jocelyne DiRuggiero

**Author notes:** Corresponding author: Jocelyne DiRuggiero Johns Hopkins University Biology Department 3400 N. Charles Street, Mudd Hall Baltimore MD 21218, USA.

## Abstract

Halite endoliths in the Atacama Desert represent one of the most extreme microbial ecosystems on Earth. Here we sequenced and characterized a shotgun metagenome from halite nodules collected in Salar Grande, Chile. The community is dominated by archaea and functional analysis attributed most of the autotrophic CO_2_ fixation to a unique cyanobacterium. The assembled 1.1 Mbp genome of a novel nanohaloarchaeon, *Candidatus* Nanopetramus SG9, revealed a photoheterotrophic life style and a low median isoelectric point (pI) for all predicted proteins, suggesting a “salt-in” strategy for osmotic balance. Predicted proteins of the algae identified in the community also had pI distributions similar to “salt-in” strategists. The Nanopetramus genome contained a unique CRISPR/Cas system with a spacer that matched a partial viral genome from the metagenome. A combination of reference-independent methods identified over 30 complete or near complete viral or proviral genomes with diverse genome structure, genome size, gene content, and hosts. Putative hosts included *Halobacteriaceae*, *Nanohaloarchaea*, and *Cyanobacteria*. Despite the dependence of the halite community on deliquescence for liquid water availability, this study exposed an ecosystem spanning three phylogenetic domains, containing a large diversity of viruses, and a predominant “salt-in” strategy to balance the high osmotic pressure of the environment.

## Introduction

In the most arid deserts on Earth, microorganisms find refuge inside rock substrates as a survival strategy (Pointing and Belnap, 2012, Wierzchos *et al*., 2012b). The rock environment provides physical stability, protection from incident UV and excessive shifts in temperature, and enhances moisture availability (Chan *et al*., 2012, Walker and Pace, 2007). The colonized substrates are translucent, allowing primary production to occur via photosynthesis (Walker and Pace, 2007, Wierzchos *et al*., 2012b). These endolithic communities are typically composed of cyanobacteria associated with diverse heterotrophic bacteria and/or archaea, and sometimes eukaryotes (Chan *et al*., 2013, Robinson *et al*., 2015, Wierzchos *et al*., 2012b). The diversity of rock habitats colonized by microorganisms has shown that life has found innovative ways to adapt to the extreme conditions of hyper-arid deserts (Friedmann 1982, DiRuggiero *et al*., 2013, Pointing *et al*., 2009, Wei *et al*., 2015, Wierzchos *et al*., 2012b). This is in stark contrast to soil, where microorganisms under extreme water stress and restricted access to nutrient must undergo long periods of stasis (Crits-Christoph *et al*., 2013).

The Atacama Desert in Northern Chile is one of the oldest and driest deserts on Earth (Clarke 2006). In the hyper-arid zone of the desert, with decades between rainfall events and extremely low air relative humidity (RH) (mean yr^-1^ values <35%), deliquescence of ancient halite crusts of evaporitic origin was shown to provide sufficient moisture to sustain microbial communities (Davila *et al*., 2008, de los Rios *et al*., 2010, Robinson *et al*., 2015, Wierzchos *et al*., 2012a). Within the halite nodules, capillary condensation of water vapor at air RH as low as 50-55%, due to the presence of pores smaller than 100 nm surrounding large NaCl crystals inside the nodules, was reported as a potential source of water for microorganisms (Davila *et al*., 2008, Wierzchos *et al*., 2012a). Under these conditions, the halite nodule interior in the Yungay area of the hyper-arid core remained wet for 5,362 hours yr^−1^ (Wierzchos *et al*., 2012a). In contrast, in Salar Grande, located in the southwest area of the Tarapacá Region, coastal fogs are frequent (Cereceda *et al*., 2008a, Cereceda *et al*., 2008b) leading to constant moisture inside the nodules (Robinson *et al*., 2015).

High-throughput culture-independent methods based on 16S rRNA gene sequencing have shown that the Atacama halite communities were dominated by archaea from the *Halobacteriaceae* family. The communities also contained a unique cyanobacterium related to *Halothece* species and diverse heterotrophic bacteria (de los Rios *et al*., 2010, Robinson *et al*., 2015). Halite communities exposed to costal fogs were more diverse and harbored a novel type of algae that was not found in the Yungay nodules, suggesting that the environmental conditions in this habitat might be too extreme for eukaryotic photosynthetic life (Robinson *et al*., 2015).

Assimilation of atmospheric radiocarbon into the halite microbial community biomass showed that carbon cycling inside the halite nodules was ongoing, with carbon turnover times of less then a decade in Salar Grande (Ziolkowski *et al*., 2013). Measurements of chlorophyll fluorescence using Pulse Amplitude Modulated (PAM) fluorometry recently demonstrated *in situ* active metabolism in halite endolithic communities (Davila et al., 2015). Photosynthetic activity was tightly linked to moisture availability and solar insolation and was sustained for days after a wetting event (Davila *et al*., 2015). Radiolabelled experiments showed that the halite communities fixed CO_2_ via photosynthesis and further evidence of metabolic activity was supported by oxygen production and respiration (Davila *et al*., 2015).

To further characterize halite endoliths, we sequenced the pooled metagenome of a microbial community associated with halite nodules. We found novel microorganisms, community members from the three domains of life, and a large diversity of viruses. The functional annotation of the metagenome revealed communities highly specialized to the extreme salinity of the environment.

## Materials and methods

### Sampling, DNA extraction, and sequencing

The colonization zone from 5 halite nodules collected in Salar Grande (Fig. 1) was harvested using aseptic conditions in a laminar flow hood and pooled together for DNA extraction, as previously described (Robinson *et al*., 2015). Sequencing was performed on the Illumina HiSeq2500 at the University of Maryland School of Medicine Institute for Genome Sciences (Baltimore, MD).

**Figure 1:**
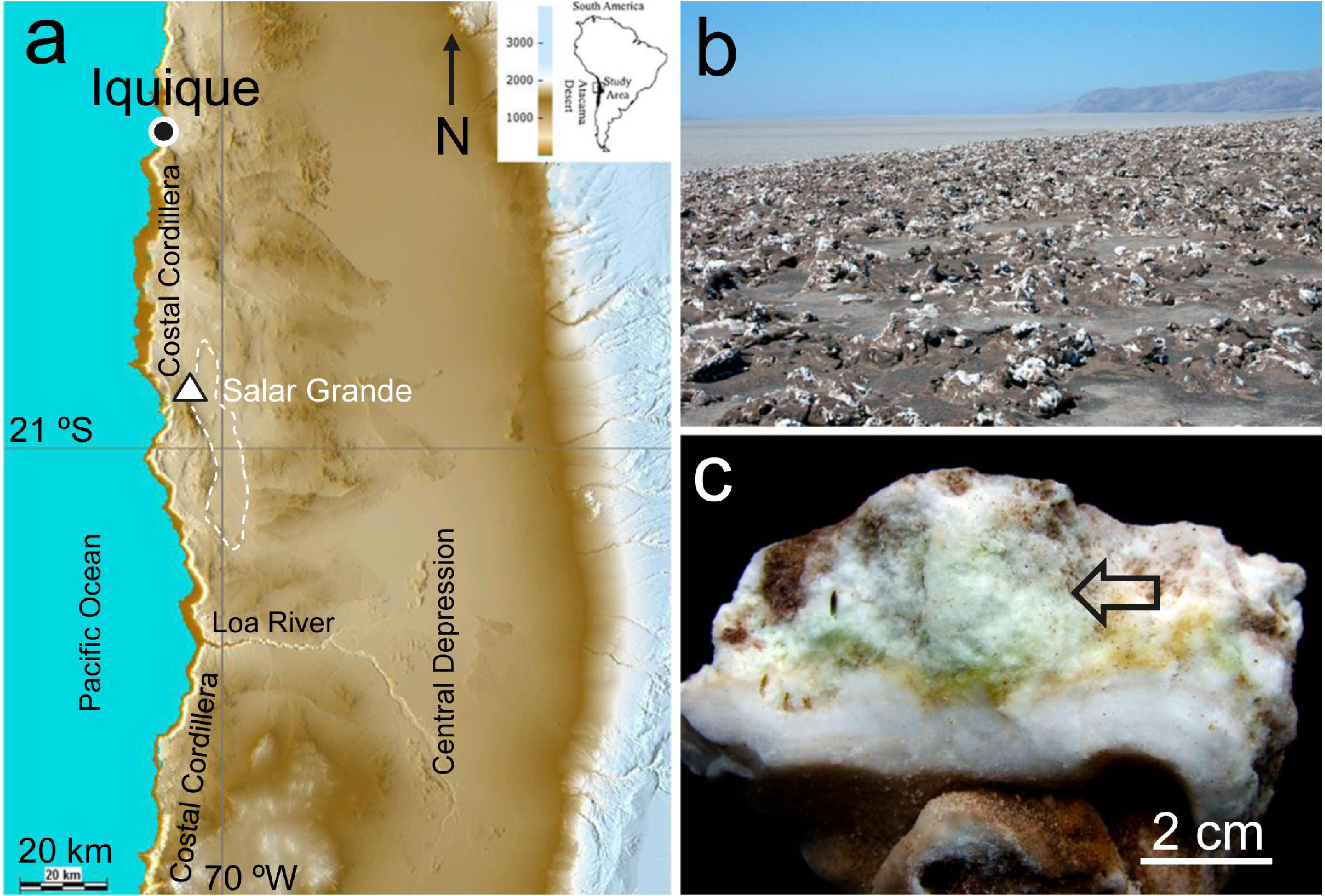
(a) Shaded relief digital map of the northern Atacama Desert, Chile, with the Salar Grande sampling location (triangle); (b) Salar Grande halite nodule field; (c) Section of a halite nodule with a back arrow indicating the green diffuse colonization zone.

### Whole Metagenome Analysis

Sequencing produced 95,230,365 paired-end reads. After quality control (see details in supplementary material), reads were assigned taxonomy content using PhyloSift (Darling *et al*., 2014) and the functional content was characterized using the analysis server MG-RAST (Meyer *et al*., 2008). The metagenome was assembled using the IDBA-UD assembler (Peng *et al*., 2012). We use a *k* range of 20-100 with a pre-correction step before assembly, producing a meta-assembly with a mean contig size of 1,060 bp, a max contig size of 377,822 bp, and a contig n50 of 1,558 bp. Assembled contigs were grouped into potential draft genomes using tetranucleotide frequencies, abundance levels, and single-copy gene analysis with MaxBin (Wu *et al*., 2014). Genomic bins were assigned taxonomic ranks with PhyloSift and Kraken (Darling *et al*., 2014, Wood and Salzberg 2014).

### Algae genome

Genomic bin 18 contained a high number of eukaryotic marker genes (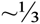 of all marker genes in the bin), all of which were identified to belong to a member of the eukaryotic green algae using BLASTP against the non-redundant (nr) database at NCBI. Contigs assembly and annotation are further described in supplementary material.

### Nanohaloarchaea genome

Four of the largest assembled contigs were binned together with MaxBin (Wu *et al*., 2014) and identified as the partial genome of a member of the Nanohaloarchaea. Reassembly methods (see details in supplementary material) resulted in a single assembled genomic contig. The completed assembly was uploaded to and annotated using the RAST server (Aziz *et al*., 2008). A CRISPR/Cas system was annotated using RAST and CRISPRFinder (Grissa *et al*., 2007). BLASTN was used to match CRISPR spacers to the assembled metagenome content.

Phylogenetic positioning of the novel *Candidatus* Nanopetramus SG9 genome was performed by extracting 12 ribosomal marker genes shared by archaeal genomes with PhyloSift (Darling et al 2014), aligning with MUSCLE (Edgar, 2004), and building a Maximum-Likelihood tree with FastTree (Price *et al*., 2010) using concatenated conserved blocks from each alignment (Gblocks) (Castresana 2000). The G+C content of the CRISPR/Cas system was calculated and compared to the average G+C content for 10 kbp windows across the entire genome. The phylogenetic position of the CRISPR/Cas system was determined by aligning Cas1 proteins from key archaeal species using MUSCLE and by building a Maximum-Likelihood tree using FastTree.

### Viral genomes

VirSorter (Roux *et al*., 2015) was used to extract viral genomic content from the assembled metagenome and viral contigs greater than 12 kbp were annotated and examined. All contigs greater than 5 kbp were checked for evidence of circularity using a custom Python script, CircMG (https://github.com/alexcritschristoph/CircMG). The RAST annotation server failed to annotate the majority of proteins encoded on the viral contigs. ORFs were predicted for each putative genome with Prodigal (Hyatt *et al*., 2010). To compare relationships within the halite viral community, all predicted viral proteins were compared against all others using BLASTP and an e-value cutoff of 0.001, producing a protein-protein similarity network. This network was used to build a virus-virus weighted undirected network, where the edge weight between two viruses was determined by the sum of the amino acid percent identity of all protein-protein matches between two viruses, divided by the total number of predicted genes in both viral genomes.

Community finding was run on the virus-virus network using the Walktrap community finding algorithm (Pons and Latapy. 2005). All network analyses were done in R using the igraph package (Csardi and Nepusz, 2006). The largest clusters in the protein-protein network (representing conserved proteins) were annotated using BLASTP to the nr protein database (Gish and States 1993) and HHPred (Söding *et al*., 2005).

### Sequence data and availability

All sequences were deposited at the National Center for Biotechnology Information Sequence Read Archive under Bioproject PRJNA296403 and accession number SRP064713. The MG-RAST report for the data is available under ID 4600831.3 (http://metagenomics.anl.gov/linkin.cgi?metagenome=4600831.3).

Completed assemblies, annotation, and phylogenetic trees are available at http://figshare.com/s/02565916783811e58e4b06ec4bbcf141

## Results

### Halite metagenome taxonomic and functional analyses

We characterized at the molecular level the endolithic microbial community from halite nodules from Salar Grande (Fig. 1). Due to the difficulty in harvesting enough DNA from the colonization zone of a single halite nodule, the metagenome was obtained with DNA extracted from 5 different nodules. The metagenome sequence of the halite community was composed of 9.6 Gb of high quality, paired-end, metagenomic shotgun sequences. Taxonomic assignments of the metagenomic reads performed with PhyloSift (Darling *et al*. 2014) revealed a community dominated by Archaea (71%) and also composed of Bacteria (27%) and Eukarya (1%) (Fig. 2). *Halobacteria* represented the majority of the Archaea (90%) with a small representation of *Nanohaloarchaea* (~2%). Most bacteria belonged to the *Salinibacter* genera (63%) and cyanobacteria constituted 15% of the bacteria. Reconstruction of 16S rRNA gene sequences from the metagenomic dataset with EMIRGE (Miller *et al*., 2011) provided full-length genes for all the major taxonomic groups and was consistent with our previous work using 16S rRNA gene sequencing (Robinson *et al*., 2015) (Fig. S1).

**Figure 2:**
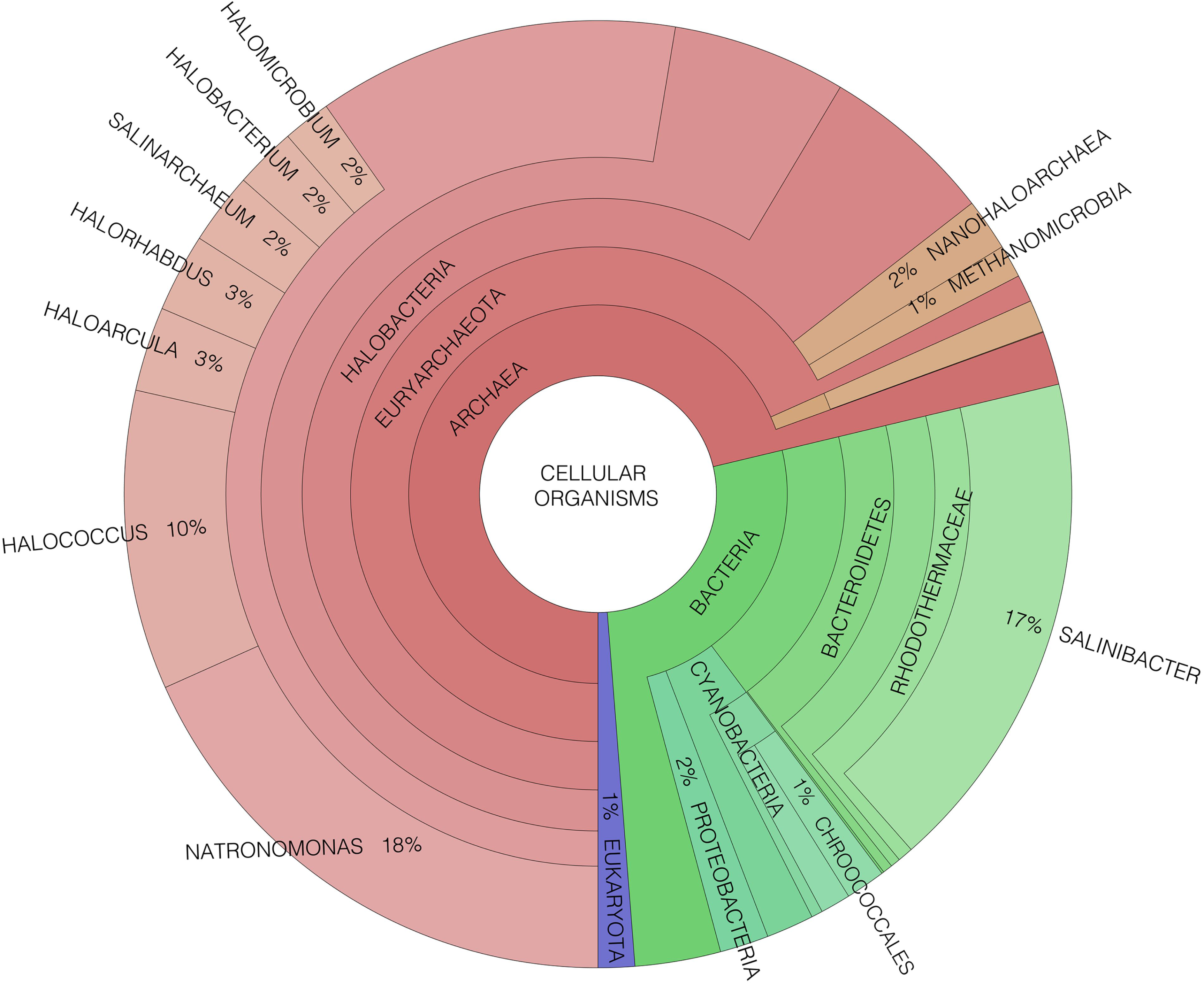
Taxonomic assignments of the halite metagenome sequence reads using Phylosift and displayed with Krona.

The functional composition of the halite metagenome was analyzed with MG-RAST (Meyer *et al*., 2008) using total sequence reads (Fig. S2). Of the genes involved in carbon metabolism, only 8% were allocated to autotrophic CO_2_ fixation (Fig. S3a) and the Calvin-Benson cycle (CB) was the only pathway for autotrophic CO2 fixation (Fig. 3b). The majority of the assigned RubisCO type I genes (>91%) were attributed to members of the *Cyanobacteria* (Fig. S4). We identified a small number of RubisCO type III genes and those were all from members of the *Halobacteriaceae*. Other key enzymes of the CB pathway, phosphoribulokinase and sedoheptulose-1,7-bisphosphatase, were also present and more than 99% of those sequences belonged to *Cyanobacteria*. Pathways for CO_2_ concentration (carboxysomes) and phosphoglycolate detoxification (photorespiration) were also present in the halite metagenome and were from *Cyanobacteria* (Fig. S3b).

**Figure 3:**
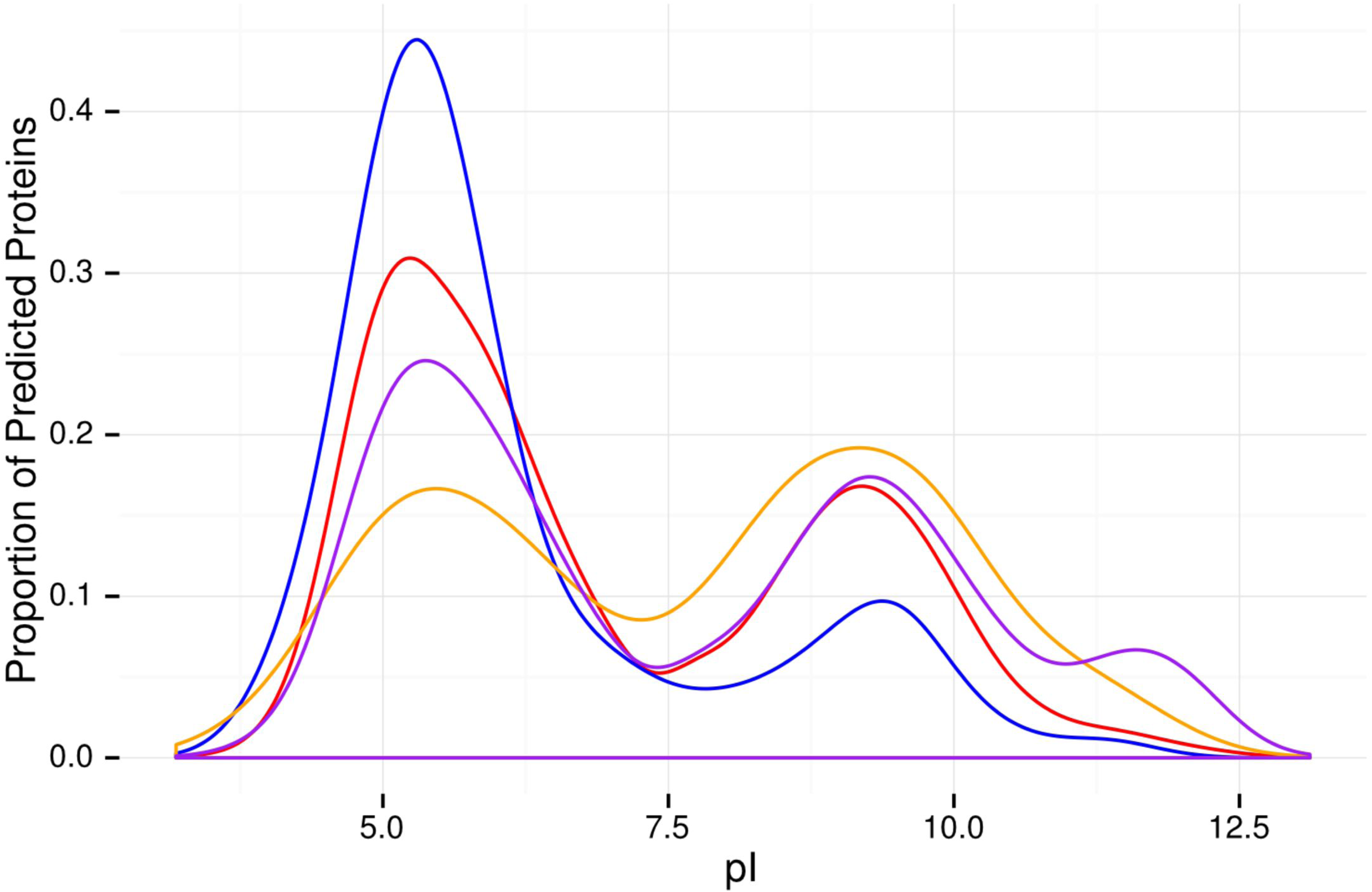
Comparison of isoelectric point profiles of the predicted proteomes for the halite alga (blue) and 3 closely related halophilic algae, *Micromonas sp. RCC299 (red), Ostreococcus tauri (purple),* and *Dunaliella salina (orange).* All reported protein sequences in the nr database were used for *Micromonas sp. RCC299* and *O. tauri.* Sequences from the UniProt Protein database were used for *D. salina.*

With respect to photosynthesis, most of the genes for Photosystem I and II (PSI and PSII) major proteins, and for light harvesting complexes, belonged to *Cyanobacteria* with only a small fraction assigned to green algae (Fig. S5 and S6). For all photosystems, 24 to 26% of all sequences were not given a taxonomic rank and 3% of PSII sequences were assigned to unclassified viruses (Fig. S5b). Phototrophy was also supported via light-driven proton pumps that belong to a number of heterotrophs including *Roseiflexus* species (proteorhodopsin), *Salinibacter* (xanthorhodopsin), and *Halobacteriaceae* (bacteriorhodopsin) (Fig. S7). Surprisingly, no nitrogenase *(nif)* genes were detected (Fig. S8).

### Osmotic adaptation of the algae from the halite community

We found 89 complete or partially complete genes identified as belonging to a eukaryotic alga with an average coverage of 7.2x and a maximum contig length of 3.9 kbp. The majority of these genes mapped closely to homologs in either *Ostreococcus tauri* or *Micromonas* sp. RCC299. Known genes included heat shock protein 70, DNA mismatch repair protein MSH4, and multiple RNA splicing factors. Eukaryotic translation initiation factors, RNA polymerase subunits, and multiple enzymes and ribosomal proteins were also identified (Table S2). Using concatenated sequences of chloroplast and mitochondrial genes, we found that the alga clustered with other members of the *Chlorophyta* (Fig. S9), grouping consistently with the *Micromonas and Ostreococcus* species and thereby confirming our previous phylogenetic position (Robinson *et al*., 2015).

To reveal potential adaptations to high salt, we compared the isoelectric point (pI) of the halite alga predicted proteins with that of the translated proteomes for *Micromonas* sp. RCC299, *O. tauri,* and the reported proteins for *Dunaliella salina,* all belonging to halophilic algae (Fig. 3) (Paul *et al*., 2008). Proteins from the halite alga had a statistically significantly lower mean pI (6.19) than the known reference proteins from *Micromonas* sp. RCC299 (6.97), *O. tauri* (7.60), and *D. salina* (7.7) (Mann-Whitney t-test; p<0.001). Isoelectric point distributions were compared using nonparametric statistical tests. Using a paired one-sided Wilcoxon t-test, we found that the predicted halite alga proteins had significantly lower pI when paired using BLASTP with *O. tauri* homologues (n=85; p<0.001; difference of means: 0.55) and with *Micromonas* sp. RCC299 homologues (n=87;p=0.025; difference of means: 0.24). A paired-sample bayesian model comparison, implemented with BEST (Bååth, 2014, Kruschke, 2013), reported that the probability that the proteins of the halite alga had a lower mean pI (difference of means less than 0) than that of *Micromonas* sp. RCC299 was 97.9%. This analysis predicted that the halite alga might have one of the lowest protein pI distributions of any reported eukaryote.

The halite metagenome also contained a number of genes from algal organelles. A 29.8 kbp contig with 39% G+C carried several chloroplast genes with high similarity to the *O. tauri* chloroplast genome, which is 71.7 kbp and 39.9% G+C. Genes encoding for PSI and PSII protein subunits, cytochrome protein subunits, ribosomal proteins, and for rRNAs and tRNAs were also found on the 29.8 kbp contig. In the same genomic bin, we also found 8 non-overlapping contigs that contained mitochondrial genes from algae. The combined non-overlapping contigs were 45.4 kbp in length with 37.1% G+C, similar to the 44.2 kbp mitochondrial genome of *O. tauri* (38.2% G+C). These genomic fragments were found to be enriched in tRNAs and organelle genes. Predicted proteins for the organelles had mean pI values above 8, which may indicate different environmental conditions in the organelle than in the intracellular space (Table S2).

### The complete genome sequence for a novel Nanohaloarchaea

Our genome assembly produced a nanohaloarchaeon genome of 1.1 Mbp long, encoding for 1,292 genes, and with a G+C content of 46.4% (referred to as SG9) (Fig. 4). Although read abundances show the assembled contigs represented only ~1-3% of the population, these contigs likely assembled well because of the small genome size (1.1 Mbp), low levels of micro-diversity, and a genome coverage around 20. A phylogenetic analysis, using a set of 12 conserved concatenated genes, showed that *Candidatus* Haloredivivus (Ghai *et al*., 2011) was the closest known reference (Fig. 5). The 16S rRNA gene sequence of SG9 was 91% identical to that of *Candidatus* Haloredivivus, and 90 and 88 % identical to *Candidatus* Nanosalina and *Candidatus* Nanosalinarum, respectively, and in agreement with our concatenated protein phylogeny. We have named this new microorganism, SG9, as *Candidatus* Nanopetramus SG9 (petramus: rock).

**Figure 4:**
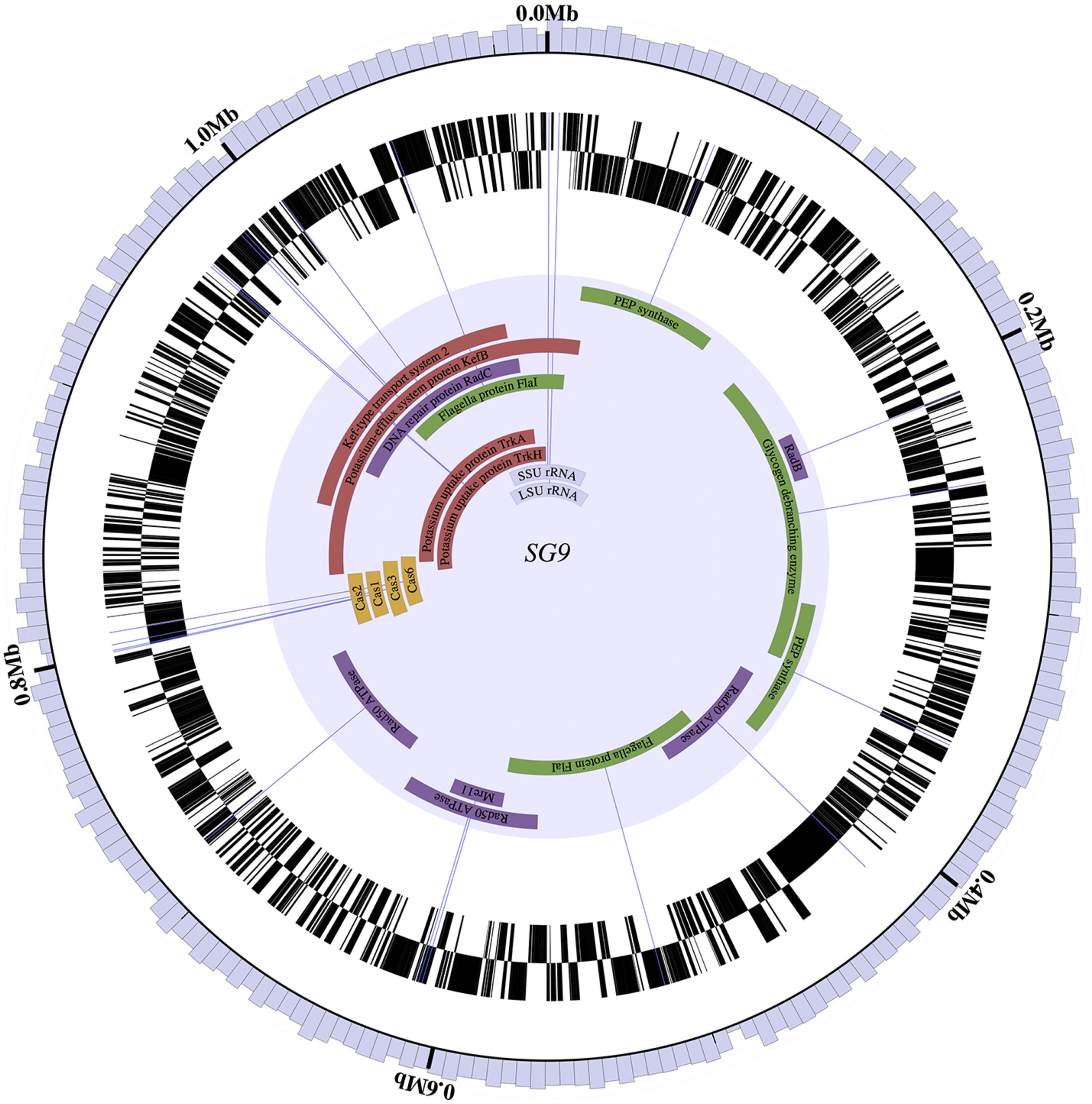
Circular representation of the SG9 genome using the Circleator tool (Crabtree *et al*., 2014). The G+C% of a 10 kbp window is displayed on the outermost circle (G+C scale: 40 to 50%). Following circles represent predicted genes on the forward strand and reverse strands, respectively; genes related to potassium homeostasis and uptake are in red, genes related to a heterotrophic lifestyle are in green, genes related to DNA repair are in purple, and Cas genes are in orange. The position of the ribosomal rRNA genes is indicated in grey.

**Figure 5:**
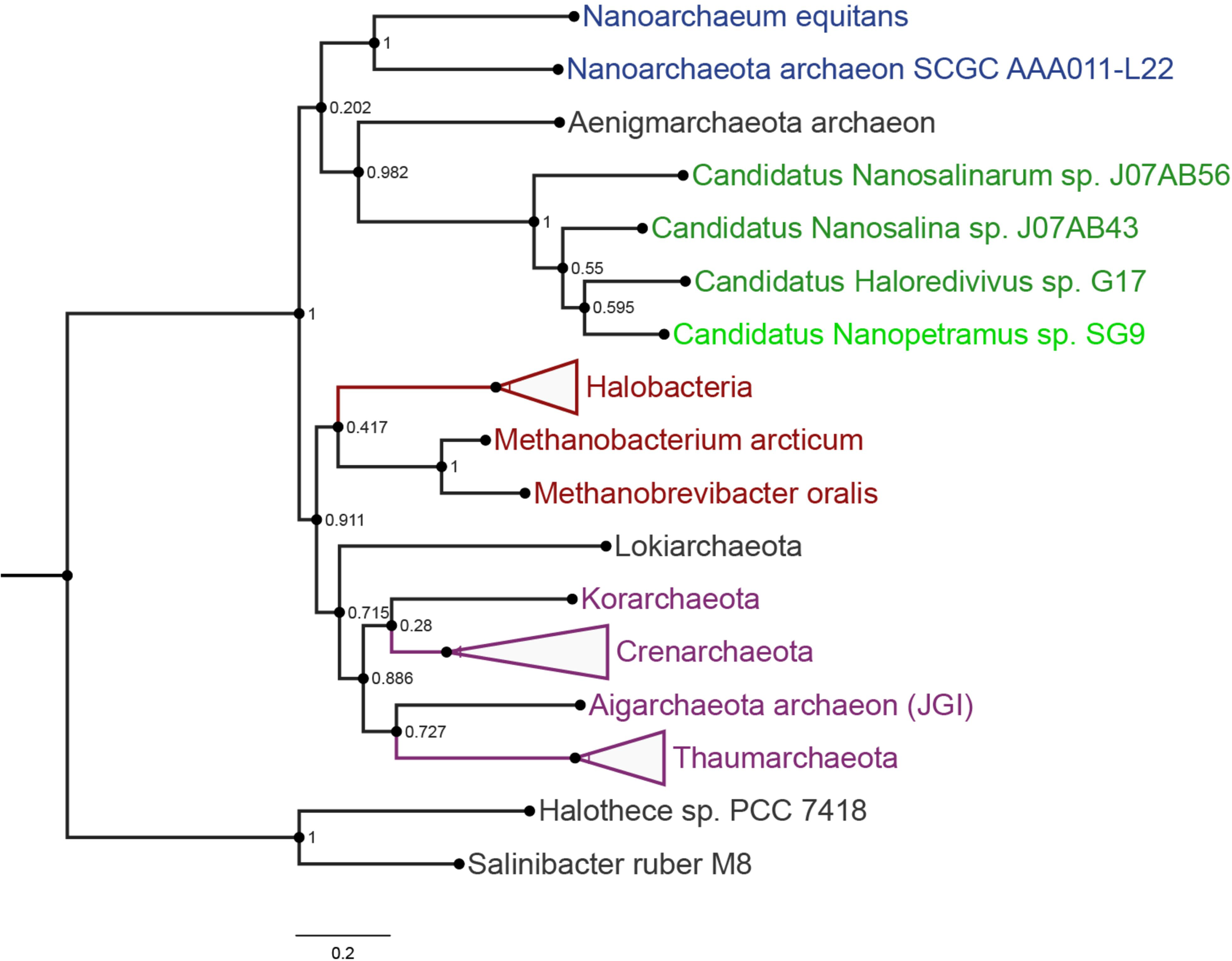
Phylogenetic position of the novel *Candidatus* Nanopetramus SG9 genome within the Archaea. The tree was built with alignments of concatenated genes for *rpsB*, *rplA, IF-2, rpsI, S5, S7, rplF, rplE, rpsK, S8, L18P/L5E,* and *rplM.* Euryarcheota are in Red, the TACK phyla in purple, Nanoarchaeota in blue, and the Nanohaloarcheota in green. Bacterial species were used as an outgroup. The scale bar represents 0.2 % sequence divergence. Bootstrap values (1000 replicates) are shown at nodes.

The SG9 genome was highly reduced and mostly composed of protein encoding genes, similar to previously reported genomes for *Nanohaloarchaea* (Narasingarao *et al*., 2012) (Fig. 4). RAST annotation of the genome revealed that 79% of predicted proteins could not be assigned to known function. We predict that SG9 has a photoheterotrophic life style as indicated by the presence of genes for rhodopsin biosynthesis, archaeal genes for carbohydrate metabolism and a phosphoenolpyruvate-dependent sugar phosphotransferase system (PTS). The glucose-6-phosphate dehydrogenase gene, essential to the Pentose-Phosphate pathway and reported in both *Candidatus* Nanosalina and *Candidatus* Nanosalinarum, was absent (Narasingarao *et al*., 2012). The presence in the SG9 genome of three potassium uptake systems, Trk, Ktr and HKT, which in bacteria are key components of osmotic regulation and resistance to high salinity (Becker *et al*., 2014), along with systems for K homeostasis, indicated a potential “salt-in” strategy for survival under high salt. This was supported by a low median pI for all *Candidatus* Nanopetramus SG9 predicted proteins (pI 4.7) and a pI distribution similar to that of *Haloarcula hispanica,* a salt-in strategist (Fig. S10). Potential motility was indicated by the presence of genes for archaeal flagellar proteins. Genes for bacterial-like and archaeal-like nucleotide excision repair pathways were also encoded in the SG9 genome, together with a photolyase gene, and several homologs for the *radA* recombinase gene.

We also found genes for isoprenoid biosynthesis, a S-layer protein, and for DNA polymerases PolI and PolII, all genes typically found in archaea.

A Type I CRISPR/Cas system composed of eight CRISPR-associated proteins and a spacer/repeat region with 22 spacers was found in the SG9 genome (Fig. 6a). We found no evidence of a CRISPR system, or individual Cas proteins, in all publicly available nanohaloarchaea genomes using CRISPR-finder (Grissa *et al*., 2007) and by searching for the Cas1 protein with a Cas1-HMM alignment using Hmmer3 (Wheeler and Eddy 2013). The 11 kbp CRISPR region of SG9 had a significantly lower G+C content of 41.5% than the whole genome (Fig. 6c). The difference in G+C%, along with the absence of any CRISPR loci in the genomes of its nearest neighbors, would indicate that SG9 acquired its CRISPR system via horizontal gene transfer (HGT) (Ochman *et al*., 2005). To further elucidate the origins of the CRISPR/Cas system in SG9, the cas1 gene product was aligned with Cas1 proteins from diverse archaeal genomes (Fig. 6b). The resulting phylogeny showed that the Cas1 protein was rooted within the *Euryarchaeota,* making it likely that the locus was acquired from *Methanobacteria* or *Halobacteria* rather than the phylogenetically closer *Nanoarchaeota*. While *Halobacteria* genomes are characterized by high G+C%, the genomes of *Methanobacteria, Nanoarchaeota,* and that of the putative nanohaloarchaeal viruses (see below) all have low G+C% similar to the CRISPR locus of SG9.

**Figure 6:**
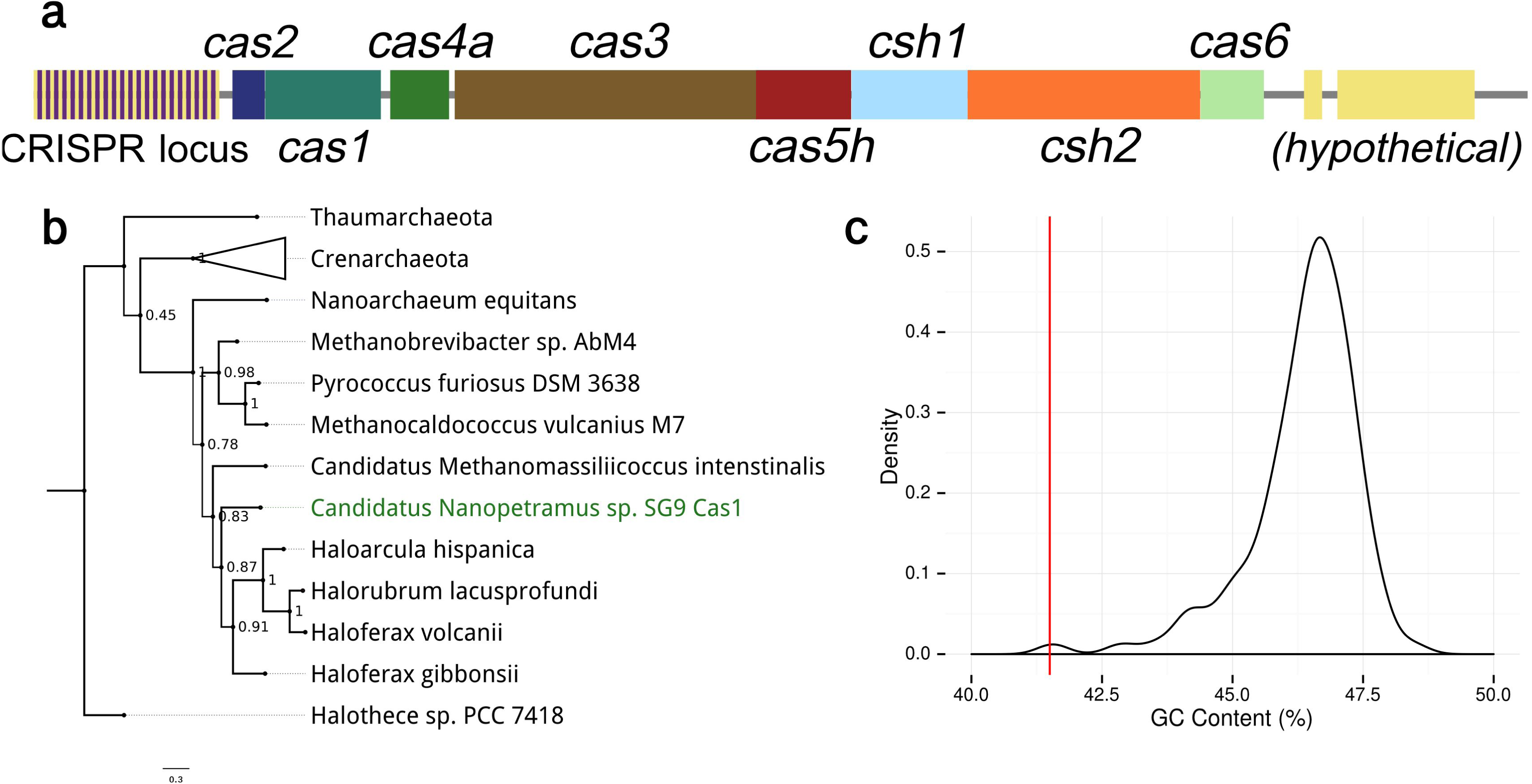
(a) A diagram of the CRISPR/Cas system found in the *Candidatus* Nanopetramus SG9 genome; (b) phylogenetic analysis of Cas1 gene products. Bootstrap values (1000 replicates) are shown at nodes; and (c) G+C% of a 10 kbp windows, with a 1 kbp step size, across the SG9 genome with the CRISPR 10 kbp region marked by a red line.

A BLASTN analysis of all 22 spacers from the SG9 CRISPR array against the assembled halite metagenome returned a single hit for spacer 22 to contig Ha1987, which was found to be a partial viral genome. Spacer 22 was located adjacent to the Cas genes cluster indicating that it was the most recently added spacer to the CRISPR array.

### Halite community viral diversity

Analysis of contigs reported by VirSorter (Roux *et al*., 2015) and of circular viral contigs (CircMG) resulted in the identification of over 30 complete or near complete viral or proviral genomes in the halite metagenome (Tables 1 and S1). These viruses are novel with a majority of viral protein products that were either hypothetical or had no homologs in the nr database. Complete genome sizes ranged from 12 kbp to 70 kbp (Table 1). While members of the *Halobacteriales* were the predicted hosts for a majority of the viruses, the *Nanohaloarchaea* were predicted as putative hosts for viral genomes Ha139 (G+C%, host genes) and Ha1987 (G+C%, spacer match), and the cyanobacterium *Halothece* for viral genomes Ha238 (host genes, G+C%) and Ha322 (host genes, G+C%), based on the evidence indicated.

**Table 1:**
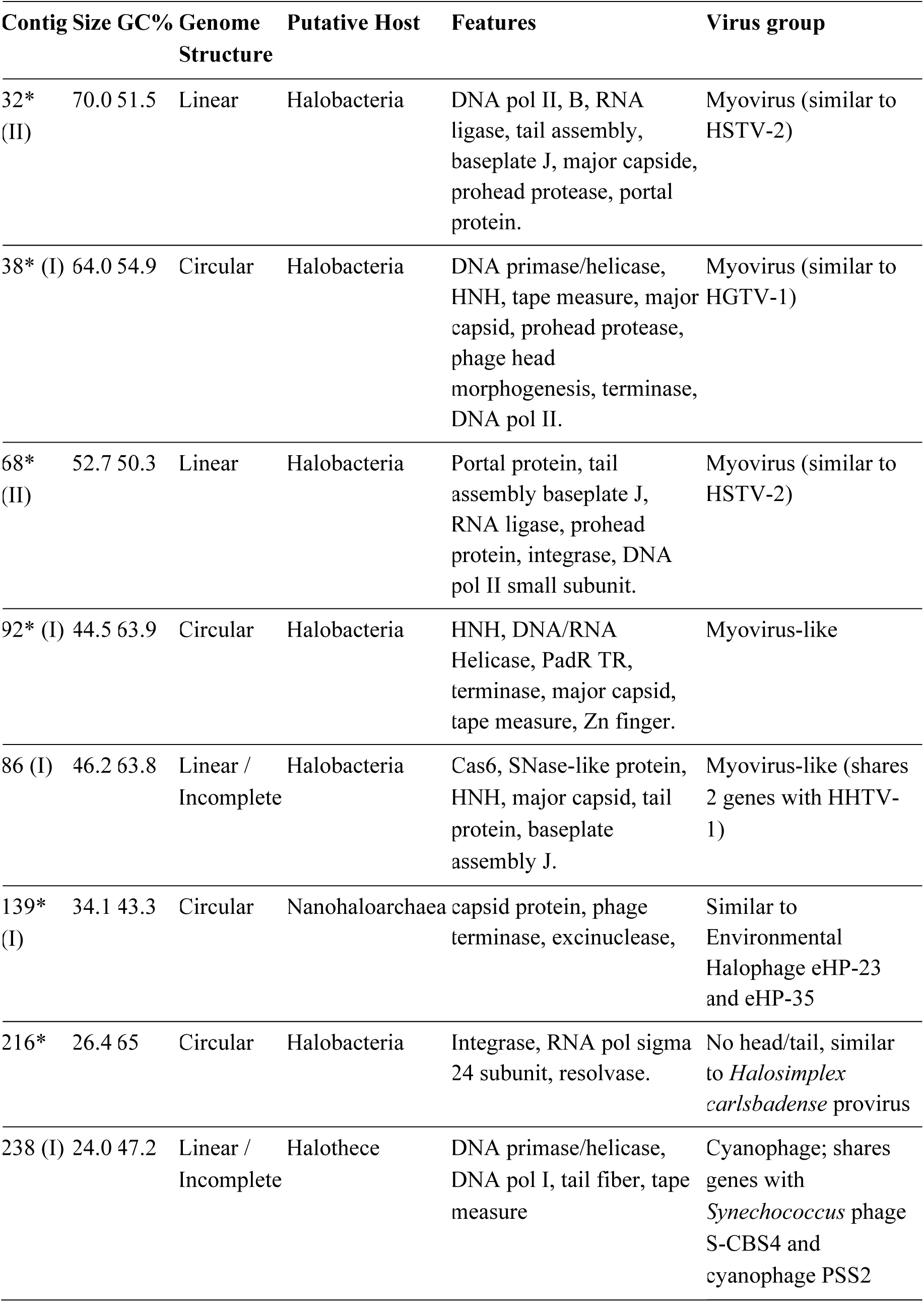
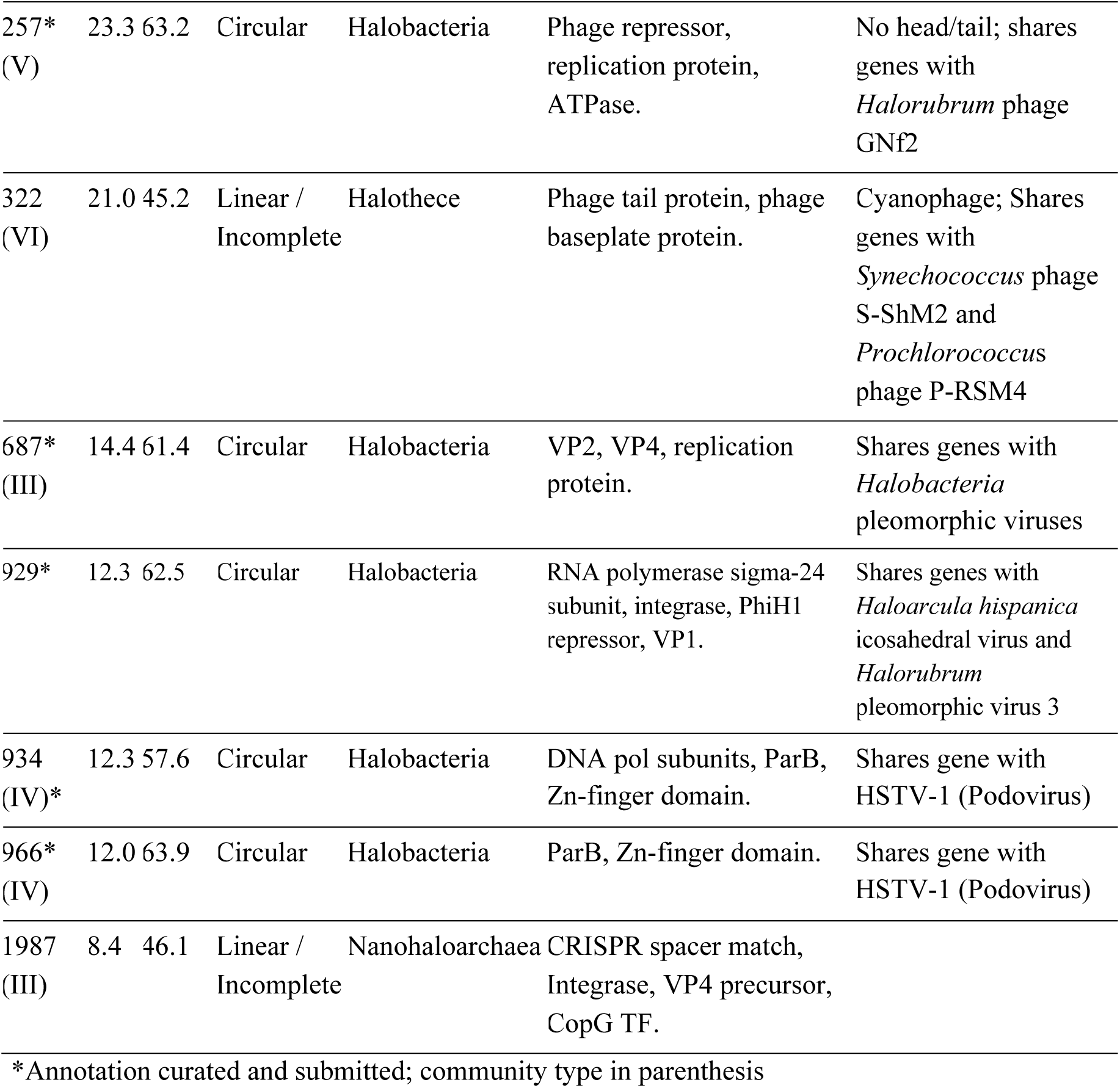
Viral genome composition and structure diversity.

A majority of the annotated viral genomes shared few to no genes with other genomes, making a phylogenetic or tree-based analysis impractical. To elucidate relationships among viruses and the structure of the viral diversity in the halite community, we used a network-based approach in which genomes were linked in a weighted network by the proportion and percent identity of predicted proteins they shared with every other genome (Fig. 7). The Walktrap community finding algorithm was used to then classify viral genomes into one of six communities (I-VI), and representative members of each cluster were further annotated and curated (Table 1)

**Figure 7:**
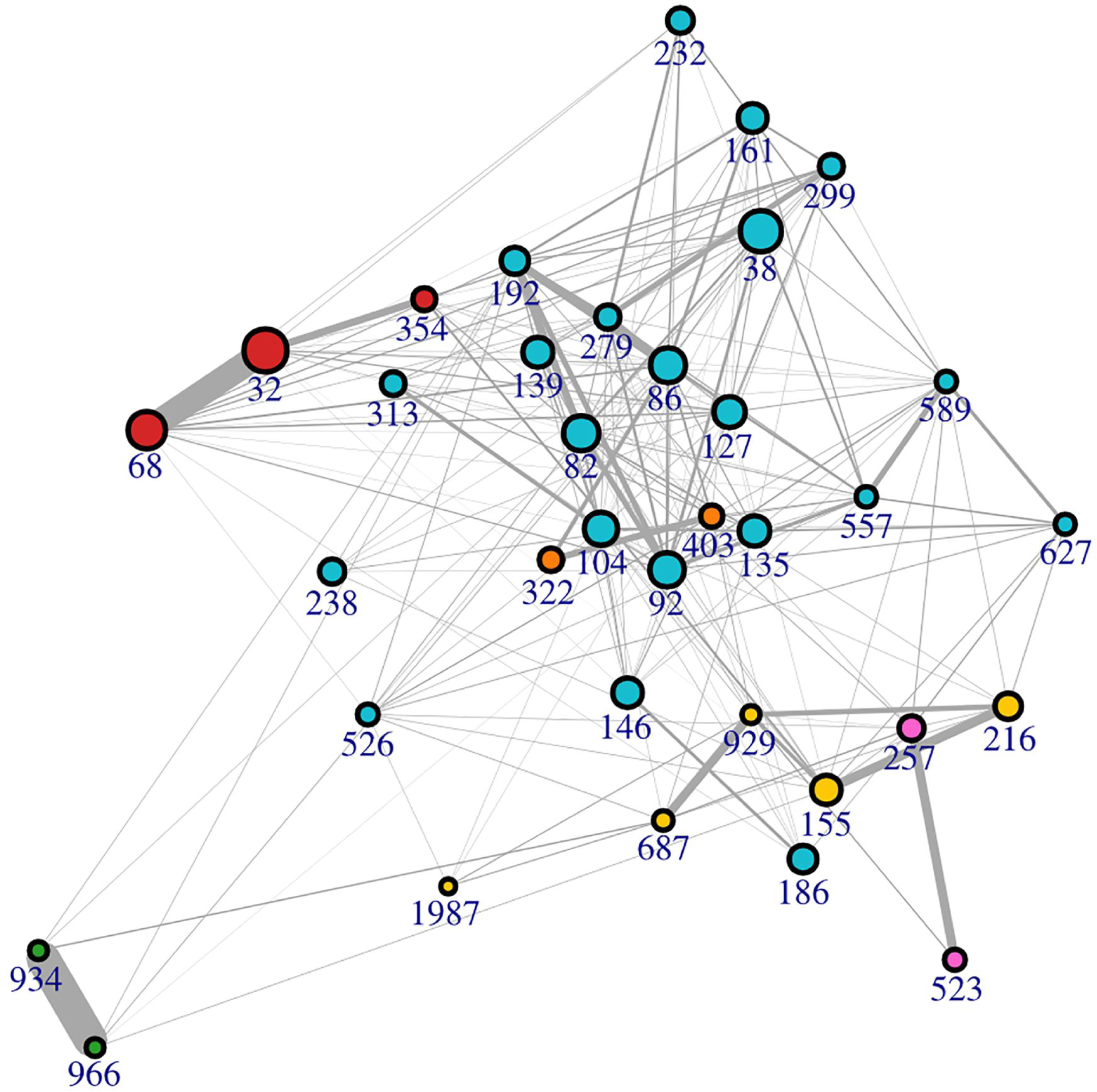
Phage-phage similarity network visualizing relationships between viral genomes. Edges are weighted based on the proportion and % ID of shared genes between two genomes. Viruses are colored according to network communities predicted by the Walktrap community finding algorithm: (I), blue; (II) red; (III) yellow; (IV) green; (V) pink; (VI) orange.

The protein similarity network used to build the network of viral relationships was examined for large clusters, which represented highly conserved proteins in the viral population. The eight largest clusters were extracted and annotated using both HHPred and BLASTP (Gish and States 1993, Söding *et al*., 2005) (Figure 8). This analysis confirmed that a majority of the viruses, particularly those in community I, had a head-tail structure. Distinct BLASTP hits to putative host transcriptional regulators were found throughout all the viral genomes and they often contained an HNH endonuclease domain. DNA polymerases were common in the larger genomes and DNA helicases were abundant in community III genomes.

**Figure 8:**
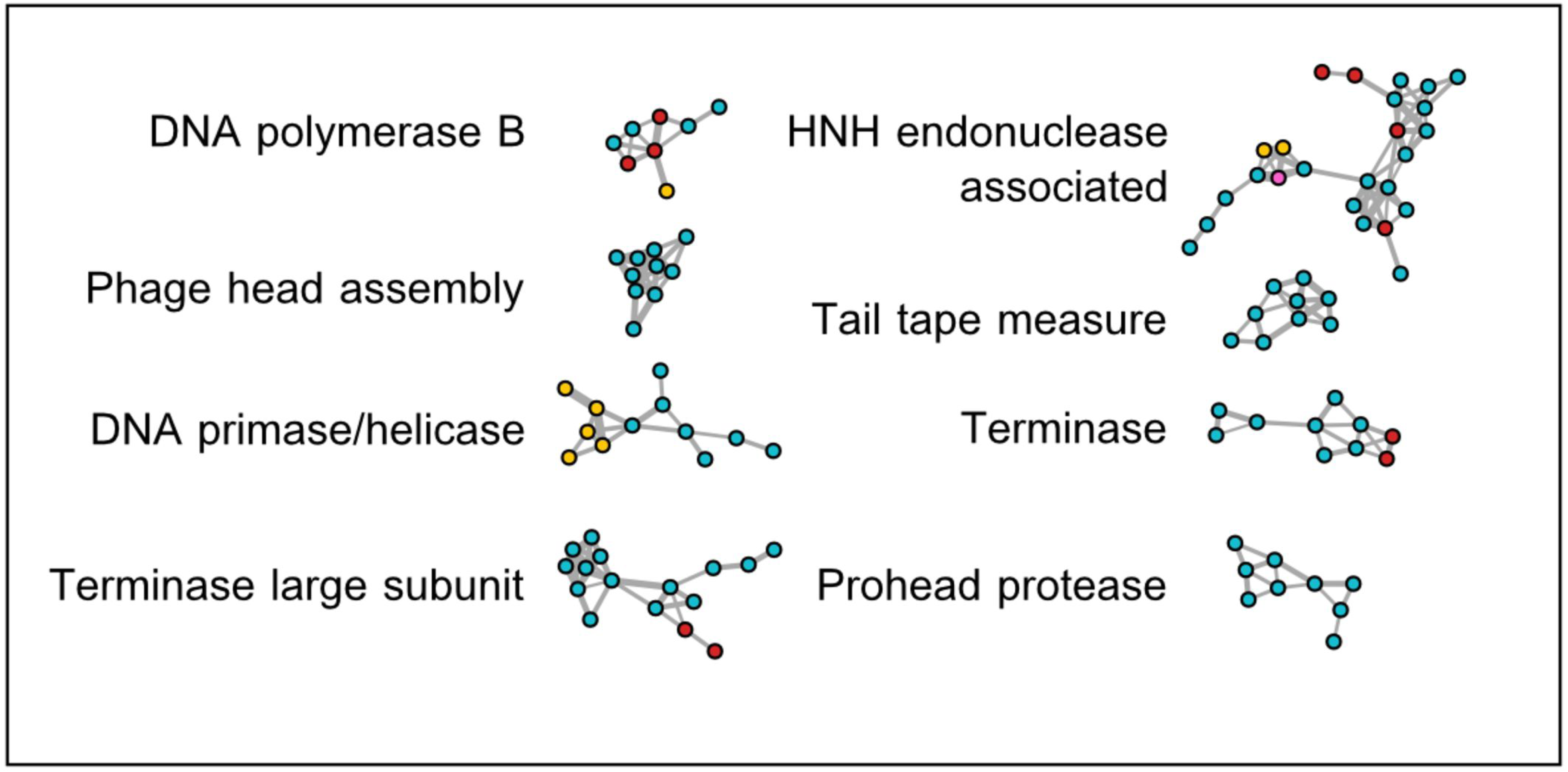
Protein-protein similarity networks of the largest clusters of viral proteins, colored by the cluster assigned to the derivative virus genome of each protein.

Among haloviruses, the largest linear genomes, Ha32 and Ha68 (community II), were structurally similar to each other and shared several genes with HSTV-2, an icosahedrally symmetric Myovirus with host *Halorubrum* (Table 1). Both contained genes for tail assembly proteins, baseplate and portal proteins, and a prohead protease, all characteristics of large head-tail viruses. The circular Ha38 (community I) genome shared few proteins with Ha32/Ha68, and most of the predicted protein products had no homologs to haloviruses in the nr protein database. Ha38 shared several core phage genes with Myovirus HGTV-1, including a major capsid protein and a prohead protease (Table 1). The genome of Ha38 also encoded DNA repair proteins (Rad25 and Hef nuclease) and a transcription initiation factor TFIIB similar to archaeal hosts. Other large, circular community I viruses (Ha86, Ha92) were not clearly related to any known halovirus and shared a small number of proteins with HGTV-1, HSTV-1, and HHTV-1, while the majority of encoded proteins had no known homologs (Table 1). Ha92 encoded for several haloarchaeal and bacterial transcriptional regulators. The Ha86 genome encoded for an ArsR family transcriptional regulator of the putative archaeal host and a *cas6* gene from a haloarchaea. Integrase genes were not found in Ha32 and Ha68, but were found in Ha216, Ha68, and Ha1987.

Genome Ha687 and the community III-associated genome Ha929 shared several genes for the structural proteins VP2 and VP4 with published *Halorubrum* pleomorphic viruses, including HRPV-1, a ssDNA halovirus (Table 1) (Pietila *et al*., 2010). Both Ha966 and Ha934 (community IV) shared a majority of their proteins with each other and few with other identified viruses from the literature or this analysis. The two genomes were both circular and approximately 12 kbp in length. They had similar structure and gene composition but surprisingly their G+C content differed by 6.6% (Table 1). Annotated proteins in their genomes included the plasmid partition protein ParB, Zn-finger domain proteins, and DNA and RNA polymerase subunits.

Genomes Ha322, Ha238, Ha1987, and Ha139 uniquely had G+C content below 48%, while haloviruses typically have GC% above 50% (Table 1) (Klein *et al*., 2002, Oren, 2006, Pagaling *et al*., 2007, Pietila *et al*., 2013b, Pietila *et al*., 2013c, Sencilo *et al*., 2013, Tang *et al*., 2002, Tang *et al*., 2004). Spacer 22 from the CRISPR-Cas array above had an exact BLAST match to the linear and partially complete genome Ha1987, linking this virus to a Nanohaloarchaeal host. In addition, several of Ha1987 viral genes had close hypothetical homologs to the published *Candidatus* Haloredivivus sp. G17 genome (Ghai *et al*., 2011). Ha139 was also putatively assigned to a Nanohaloarchaeal host because of its low G+C%, a shared hypothetical gene product with *Candidatus* Nanosalina, and because large number of gene products shared with environmental Halophage eHP-23 and eHP-35 and two viral genomes previously assigned a Nanohaloarchaeal host (Garcia-Heredia *et al*., 2012).

Both linear and incomplete genomes Ha322 and Ha238 had their closest BLASTP hits to multiple *Synechococcus* phage and cyanophage proteins, and had homologs for multiple genes in several cyanobacteria. These findings suggest that they might be novel cyanophages targeting the *Halothece* members of the community. Despite multiple close BLASTP hits to cyanophage proteins in public datasets, Ha322 and Ha238 contigs shared no homologous proteins with each other and did not cluster in the network analysis.

## Discussion

Halite nodules in fossil continental evaporites from the Atacama Desert are at the extreme of salt concentrations for hypersaline environments. Microorganisms inhabiting this environment must balance the osmotic pressure of their cytoplasm with that of the outside milieu. One osmotic strategy is the “salt in” strategy where ions, mainly K^+^ and Cl^−^, are accumulated in the cell and the entire intracellular enzymatic machinery is adapted to high salt (Oren, 2008). To remain soluble, the proteins of “salt-in” strategists have an increased number of acidic amino acid residues on their surface, resulting in a proteome with a low pI. This strategy is used by halophilic archaea and one extremely halophilic bacterium, *Salinibacter,* a member of the *Bacteroidetes* (Oren, 2008). Other halophilic and halotolerant microorganisms balance the high osmotic pressure of their environment by synthesizing compatible solutes, a strategy called “salt-out” (Galinski, 1995, Oren, 2008).

### Adaptation of the halite community to high salt

The phylogenetic composition of the halite community reflected the extreme salinity of its environment with Archaea greatly outnumbering Bacteria (Ghai *et al*., 2011, Podell *et al*., 2013, Robinson *et al*., 2015). In addition, most of the bacteria in the community belonged to the genus *Salinibacter*, a “salt-in” strategist (Oren 2008). We found only one cyanobacteria, *Halothece,* as previously described using high-throughput 16S rRNA gene sequencing (Robinson *et al*., 2015). Many species of cyanobacteria are adapted to high salt and they often form dense benthic mats in saline and hypersaline environments, where they are the main primary producers (Oren, 2015). However, above 25% NaCl only cyanobacteria of the *Aphanothecee-Halothece-Euhalothece* cluster have been found so far (de los Rios *et al*., 2010, Garcia-Pichel *et al*., 1998, Robinson *et al*., 2015, Wierzchos *et al*., 2006). A property of this cluster is the production of abundant extracellular polysaccharides (EPS) (de los Rios *et al*., 2010, Oren, 2015). We previously reported that, in the halite nodules, *Halothece* formed cell aggregates surrounded by a thick sheath embedded in EPS (de los Rios *et al*., 2010, Robinson *et al*., 2015). It is likely that these structural components play a significant role in the desiccation tolerance of *Halothece* and its ability for photosynthetic O_2_ evolution (Tamaru *et al*., 2005).

We found evidence of autotrophic CO_2_ fixation via the CB pathway by cyanobacteria in the halite metagenome. Although we also found RubisCO type III gene in several archaea, it is not clear whether this enzyme participate in autotrophic CO_2_ fixation or in a novel AMP recycling pathway (Falb *et al*., 2008, Sato *et al*., 2007). Our findings indicate that the unique cyanobacteria is likely responsible for most of the CO_2_ fixed in the halite community. In addition, we recently reported *in situ* carbon fixation through oxygenic photosynthesis in halite nodules supporting the idea that cyanobacteria are the major primary producers in this ecosystem (Davila *et al*., 2015). A number of organisms encoded light-driven proton pumps in their genomes, carrying out photoheterotrophy and significantly increasing the energy budget from light.

We assembled the partial genes of the alga previously detected in the halite community, together with large regions of the genomes of its mitochondria and chloroplast. The pI of the alga predicted proteins was one of the lowest pI reported for any eukaryote (Kiraga *et al*., 2007). In addition, the bimodal distribution of the proteins pI was similar to that of “salt-in” strategists, suggesting that lower eukaryotes might potentially use intracellular salt as a mean to balance osmotic pressure in hypersaline environments.c

### A novel nanohaloarchaeal genome with a CRISPR array

We report here the first nanohaloarchaeon genome assembled into a single scaffolded contig from metagenome data of hypersaline environments (Ghai *et al*., 2011, Martinez-Garcia *et al*., 2014, Narasingarao *et al*., 2012, Podell *et al*., 2013). The *Candidatus* Nanopetramu SG9 genome of 1.1 Mb is very similar in size to that of previously reported Nanohaloarchaea and its genomic G+C content is intermediate in the reported range (Ghai *et al*., 2011, Narasingarao *et al*., 2012, Podell *et al*., 2013). Evidence from its genome support the idea that *Candidatus* Nanopetramus SG9 has a photoheterotrophic life-style and that it uses the “salt-in” strategy to counterbalance the high salt of its environment. A low proteome pI has also been reported for *Candidatus* Nanoredivirus, suggesting that the “salt-in” strategy might be a ubiquitous feature of Nanohaloarchaea (Ghai *et al*., 2011). A unique attribute of *Candidatus* Nanopetramus SG9 was the presence of a CRISPR array on its genome, demonstrating that adaptive immunity against viruses is also a feature of Nanohaloarchaeal genomes. This is the first report of annotated CRISPR-associated features in a nanohaloarchaeal genome and documented acquisitions of CRISPR/Cas systems via HGT in the Archaea (Godde and Bickerton, 2006, Portillo and Gonzalez, 2009, Brodt *et al*., 2011) support our hypothesis that SG9 acquired its CRISPR system via HGT.

### The viral component of the halite community

With up to 10^10^ virus-like particles per ml, hypersaline systems harbor some of the highest viral concentrations of any aquatic environments (Baxter *et al*., 2011, Boujelben *et al*., 2012). In these extreme environments, with very few eukaryotes, haloviruses are likely to play an important role in shaping community structure through predation. We have assembled over 30 complete to near complete viral genomes from a metagenome obtained from the cellular fraction of the halite samples, restricting access to viruses that were either contained inside cells at the time of sampling or associated with particle surfaces. Despite this limitation, we found great viral diversity in the halite community in terms of genome structure, genome size, G+C%, and gene composition. These viruses were novel with a majority of the viral protein products having no characterized homologs. The viral genomes were not integrated within their host genomes in the metagenome assembly, suggesting that most of the viruses were lytic rather than lysogenic, in contrast to viruses found in high temperature environments (Anderson *et al*., 2015).

Viruses infecting haloarchaea come in a variety of virion morphotypes, including spindle-shaped, pleomorphic, icosahedral, and head-and-tail (Atanasova *et al*., 2012, Pietila *et al*., 2013a, Roine and Oksanen, 2011). To date, 43 haloarchaeal tailed viruses have been reported (Atanasova *et al*., 2012, Kukkaro and Bamford, 2009, Sabet, 2012) and 17 completely sequenced genomes, comprised approximately 1.2 Mb of sequence information (Klein *et al*., 2002, Pagaling *et al*., 2007, Pietila *et al*., 2013b, Pietila *et al*., 2013c, Sencilo *et al*., 2013, Tang *et al*., 2002, Tang *et al*., 2004). Our analysis confirmed that the majority of the viruses we identified in the halite metagenome also had a head-tail structure, as previously found in other hypersaline environments (Garcia-Heredia *et al*., 2012). The viral genetic diversity uncovered here hints that a significant portion of the diversity of both head-tail and other viruses in halophilic environments still remains largely unexplored.

Our network-based approach allowed us to analyze viral relationships in the community and identify core protein encoding genes with similarity to previously described haloviruses and cyanophages (Klein *et al*., 2002, Oren, 2006, Pagaling *et al*., 2007, Pietila *et al*., 2013b, Pietila *et al*., 2013c, Sencilo *et al*., 2013, Tang *et al*., 2002, Tang *et al*., 2004). Haloviruses genomes have rather high G+C content (above 50% on average), also characteristic of haloarchaea, and the halite viruses followed that trend. The exceptions were putative cyanophages and Nanohaloarchaea viruses that exhibited a lower G+C content, consistent with previous findings (Emerson *et al*., 2012, Martinez-Garcia *et al*., 2014, Podell *et al*., 2013).

A number of partial halovirus genomes have been obtained from metagenomes from crystallizer ponds and the hypersaline Lake Tyrrell in Australia with potential hosts including *Haloquadratum walsbyi, Nanohaloarchaea,* and the bacterium *Salinibacter* (Emerson *et al*., 2012, Garcia-Heredia *et al*., 2012). Here we described new groups of viruses that prey on members of the *Halobacteriales,* the cyanobacterium *Halothece,* and on a newly described Nanohaloarchaeon, *Candidatus* Nanopetramus SG9. CRISPR sequences found in the newly described SG9 genome where also present on a partial viral genome, providing a direct connection between virus and host. However, these predictions remain to be tested (Martinez-Garcia *et al*., 2014).

Despite the extreme stress of the halite environment, this work demonstrates that this endolithic community is exquisitely adapted to the challenges of its environment. Metagenomic analysis revealed a relatively complex community with members from the three domains of life. It also revealed trophic levels, from photosynthetic primary producers to heterotrophs, viral predations, and specific physiological adaptations to the high osmotic pressure of the halite milieu. Further understanding of the major taxonomic groups and essential metabolic pathways underlying the functioning of this unique community will require a combination of field-based measurements of metabolic activity coupled with meta-transcriptomics, to capture gene expression levels for specific function and taxonomic groups.

## Acknowledgements

This work was funded by grant EXOB08-0033 from NASA and grant NSF-0918907 from the National Science foundation to JDR, by grant CGL2013-42509P from MINECO (Spain) to JW and by grant NNX12AD61G from NASA to AD. We thank Octavio Artieda for support in the field and Carmen Ascaso for valuable discussions.

